# EUAdb: a resource for COVID-19 test development

**DOI:** 10.1101/2020.07.30.228890

**Authors:** Alyssa Woronik, Henry W. Shaffer, Karin Kiontke, Jon M. Laurent, Ronald Zambrano, Jef D. Boeke, David H. A. Fitch

**Affiliations:** Department of Biology, New York University, 100 Washington Square East, New York, NY 10003; Institute for Systems Genetics and Department of Biochemistry and Molecular Pharmacology, NYU Langone Health, New York, NY 10016; Department of Ophthalmology, New York University School of Medicine, New York, NY 10017

## Abstract

Due to the sheer number of COVID-19 (coronavirus disease 2019) cases, the prevalence of asymptomatic cases and the fact that undocumented cases appear to be significant for transmission of the causal virus, SARS-CoV-2 (severe acute respiratory syndrome coronavirus 2), there is an urgent need for increased SARS-CoV-2 testing capability that is both efficient and effective^1^. In response to the growing threat of the COVID-19 pandemic in February, 2020, the FDA (US Food and Drug Administration) began issuing Emergency Use Authorizations (EUAs) to laboratories and commercial manufacturers for the development and implementation of diagnostic tests^1^. So far, the gold standard assay for SARS-CoV-2 detection is the RT-qPCR (real-time quantitative polymerase chain reaction) test^2^. However, the authorized RT-qPCR test protocols vary widely, not only in the reagents, controls, and instruments they use, but also in the SARS-CoV-2 genes they target, what results constitute a positive SARS-CoV-2 infection, and their limit of detection (LoD). The FDA has provided a web site that lists most of the tests that have been issued EUAs, along with links to the authorization letters and summary documents describing these tests^1^. However, it is very challenging to use this site to compare or replicate these tests for a variety of reasons. First, at least 12 of 18 tests for EUA submissions made prior to March 31, 2020, are not listed there. To our knowledge, no EUAs have been issued for these applications. Second, the data are not standardized and are only provided as longhand prose in the summary documents. Third, some details (e.g. primer sequences) are absent from several of the test descriptions. Fourth, for tests provided by commercial manufacturers, summary documents are completely missing. To address at least the first three issues, we have developed a database, EUAdb (EUAdb.org), that holds standardized information about EUA-issued tests and is focused on RT-qPCR diagnostic tests, or “high complexity molecular-based laboratory developed tests”^1^. By providing a standardized ontology and curated data in a relational architecture, we seek to facilitate comparability and reproducibility, with the ultimate goal of consistent, universal and high-quality testing nationwide. Here, we document the basics of the EUAdb data architecture and simple data queries. The source files can be provided to anyone who wants to modify the database for his/her own research purposes. We ask that the original source of the files be made clear and that the database not be used in its original or modified forms for commercial purposes.

## Database architecture

To construct EUAdb, FileMaker^™^ was chosen for its ease in building databases that can be freely shared online (e.g. via WebDirect^™^) without clients needing to use any tool other than a web browser. Although EUAdb is meant to be fairly straightforward and user-friendly for those with some knowledge of RT-qPCR tests, knowing a little about FileMaker will help users understand the data and generate custom queries. Fig. 1 shows how the data from the FDA site is organized into tables and how these tables are related to each other. <u>Laboratories</u> that have applied for EUAs are represented with data sufficient to identify them, including URLs to the lab web sites. Although it does not yet happen, each laboratory could potentially employ multiple tests. The <u>Tests</u> table includes data fields with URLs to documents stored on the FDA site, test-specific notes about sampling, reagents or instruments used, types of controls used, details about how results from the tests should be interpreted, etc. Several calculated fields are used for reporting (e.g. most recent EUA issue date represented in the database). “Join” tables are used to allow many-to-many relationships between tests and sampling techniques, reagents or instruments. That is, because a test can use multiple sampling techniques, reagents or instruments, and the same sampling technique, reagent or instrument can be used in multiple tests, special tables with double index fields are needed to establish these relationships (tables with “Jct...” in their names). These join tables are then linked to tables for specific <u>Sample Types</u>, <u>Nucleic Acid Extraction Kits</u>, <u>Master Mix Kits for PCRs</u>, *Instruments* used for either nucleic acid extraction or RT-qPCR, and <u>Primers</u> that may be parts of <u>Primer Kits</u> used in the PCR. These tables are also linked to a <u>Manufacturers</u> table (represented by multiple occurrences, one for each reagent or instrument table). Fields, tables and architecture may change somewhat in the future as new tests are developed.

**Figure 1.**
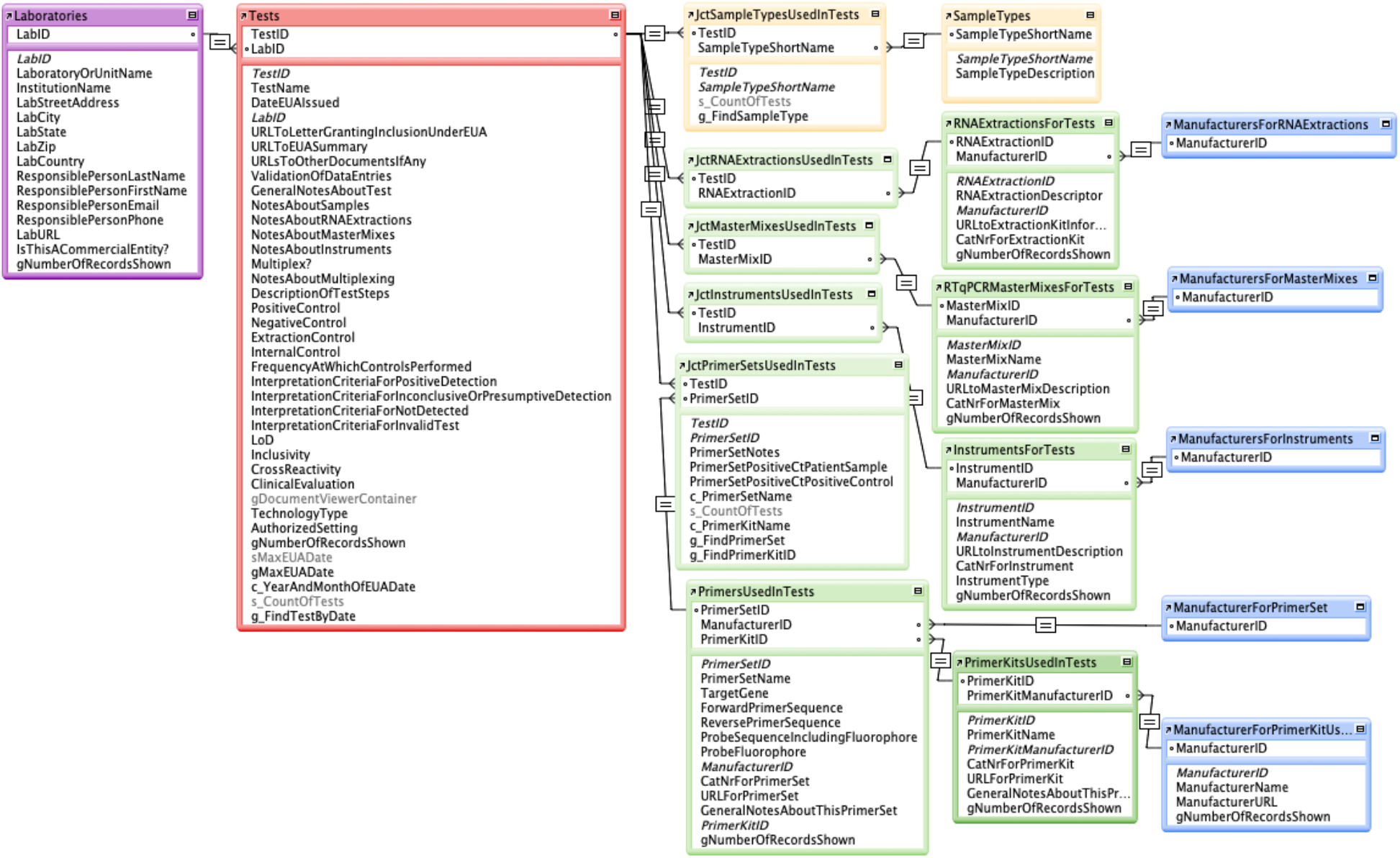
Main database architecture of EUAdb. Tables of data are related via one-to-many connections using a unique index field in each table, except for “join” tables that establish many-to-many connections via pairs of fields.

## Data curated and standardized ontology

In defining the specific data fields of the tables described above, we sought to standardize relevant data across all tests, thus allowing controlled and efficient data entry as well as comparisons among tests. However, the data provided for each test by the US FDA is in the form of a Summary document written by the laboratory developing the test. Although the FDA requests certain types of information about each test, the Summary documents are written in free-form prose and there is no easy way to automatically capture these data into individual fields. Thus, information about each test is manually transferred from the Summary document to the database by curators who are trained biologists knowledgeable about RT-qPCR testing. This curation process required deciding on specific ontology, where the same reagent, instrument, etc. may be described with slightly different terminology in different Summary documents. For example, we developed a standard set of “sample types” to describe what kinds of samples the tests were designed to use. We have also added some details that may not be explicitly found in the Summary documents, such as catalog numbers, URLs to manufacturer web sites and primer sequences, except in cases where the identity of specific reagents, instruments or manufacturers is unspecified or unclear. Data is entered by one person and independently validated by a second person. To enter data, fields show a list of values that are currently in the database and can be selected, allowing controlled data entry and efficiency as well as preventing redundancy. As new data are entered, the value lists are automatically updated.

## Accessing and using EUAdb

EUAdb is password-secured for administrative-level access, but anyone can access all the data and perform all queries via an open Guest account at EUAdb.org. This URL leads to the <u>Home</u> page (Fig. 2), which provides one-stop access to all data via global query fields and buttons attached to various scripts. In the upper left, Options-Menu and Return-to-Previous-Page buttons provide functions that are universally available on all layout pages. For example, clicking on the Options-Menu button pulls down a menu that allows you to choose the option to see a list of all tests or to exit the database. It is important to use the “Return to Previous Page”button instead of the browser “back” button because the database is run within a single browser window. A URL is provided to the US FDA site from which all information is obtained. Users are automatically signed out after 15 minutes of inactivity.

**Figure 2.**
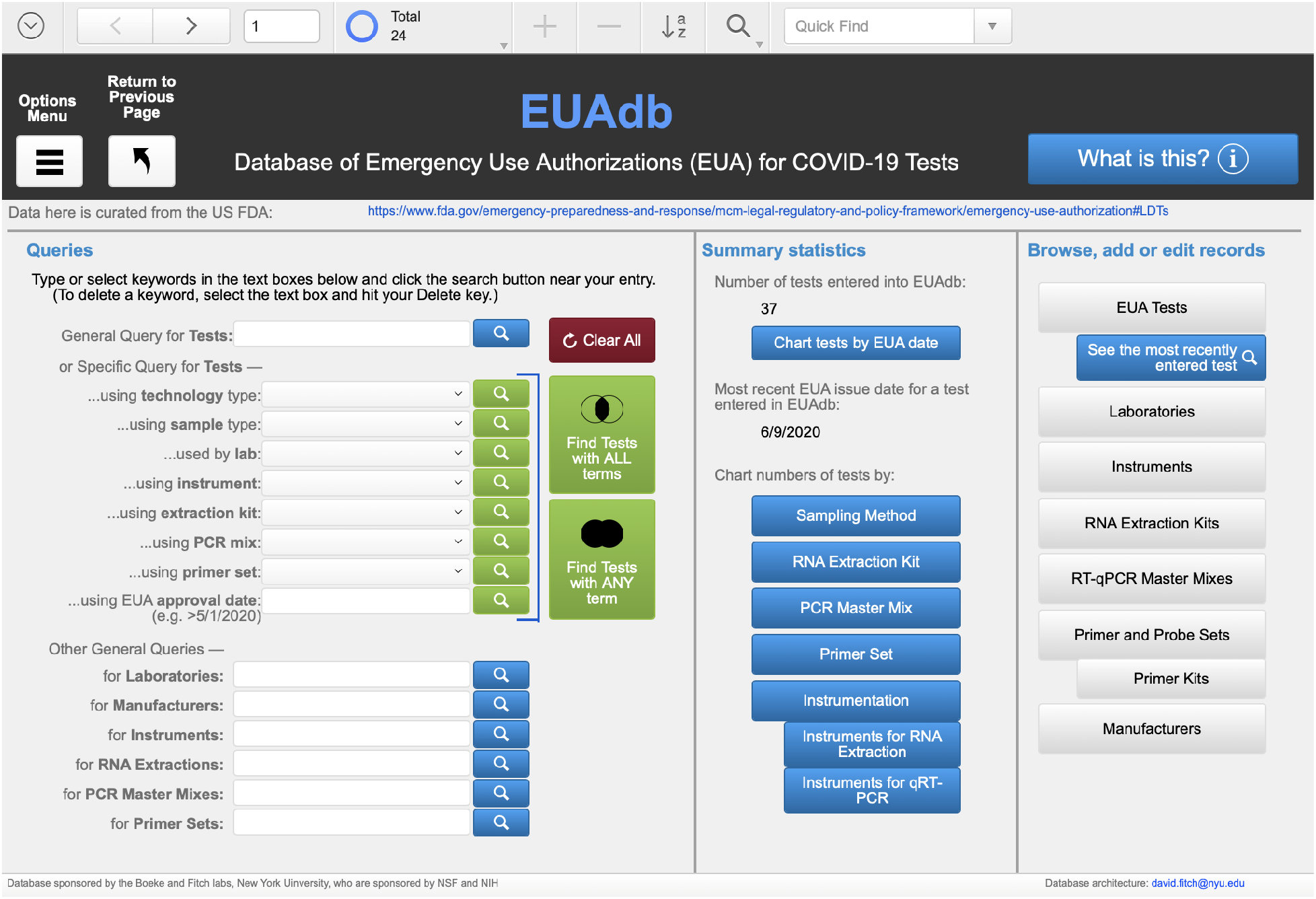
The Home layout is the landing page for EUAdb.org. See text for description.

Several types of *Queries* are available by typing keyword(s) into field(s) and selecting a button. Keyword fields next to blue query buttons allow any search term to be entered; clicking on the blue button will open a new page listing items that have the search term somewhere in the record. Several such free-form query fields are provided to query tests, labs, manufacturers, instruments and the different types of reagents. Keyword fields next to green buttons only allow particular terms to be selected in conjunction with a search for tests containing those terms. This allows a more controlled method of querying. For example, one can find all tests that use “Bronchial washes (BW)” as a sample type by selecting that term in the “Specific Query for Tests...using sample type” field and clicking the adjacent green query button. This action will bring up a page listing all tests in which bronchial washes are used as a sample type. More complex queries for tests can also be generated by selecting query terms for multiple kinds of items and using one of the large green buttons to find all tests consistent with “all” query terms (i.e., the Boolean intersection of sets of tests having each term) or with “any” of the query terms (i.e., the Boolean union of such sets). The red “Clear all” button provides an easy way to reset all terms to null values.

Several types of summary statistics are provided, including the number of tests recorded in the database, the most recent EUA issue date, and several charts about the tests and what kinds of samples, reagents or instruments are used in the tests. For example, clicking on the “Sampling Method” button brings up a page with a bar chart showing the numbers of tests that use each type of sample (Fig. 3). This chart shows that (at the time this manuscript was written) nasopharyngeal swabs are the only type of sample that is authorized by all RT-qPCR tests currently in the database and only three are recommended for saliva samples. To determine which tests these are, use the query box at the top right of the chart page to select “Saliva specimens” (you may need to select the down arrow in the value list to see additional values) and click on the green query button.

**Figure 3.**
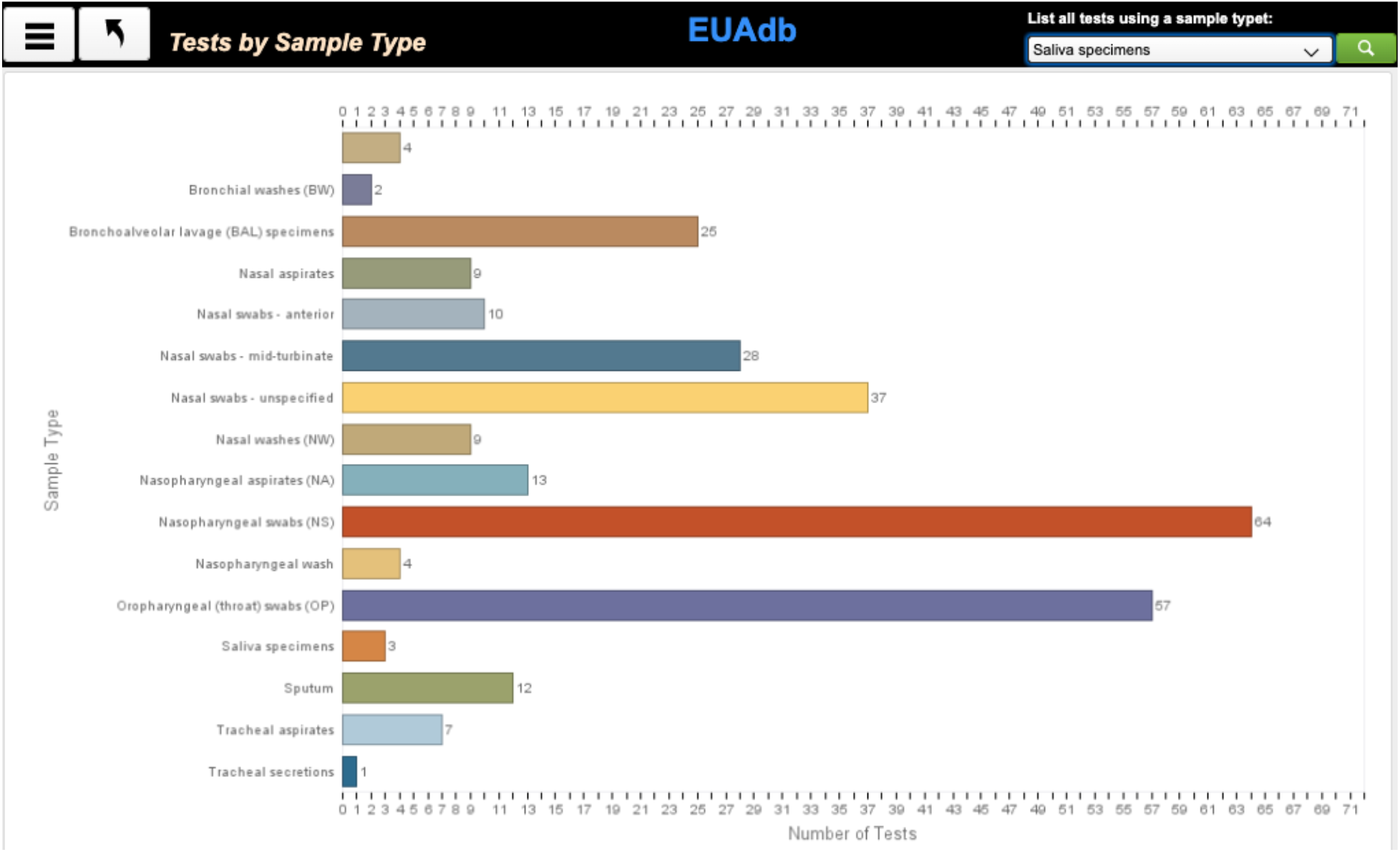
Example of a chart showing summary statistics for EUAdb, in this case the number of tests using a particular sample type. In the upper right corner is a query field for showing all tests that use a selected sample type.

On the right side of the Home page are a series of buttons that allow Guest users to browse and authorized users to add and edit all of the data in all of the tables (except the join tables). For those with FileMaker (FM) experience, typical FM functions are preserved at the top of the screen, allowing users a broader range of record navigation, query and sort functionality than provided on the Home page.

## A typical test page

Selecting “Browse all tests” from the Options menu will open a page listing all EUA tests entered into the database (Fig. 4). This only provides a few fields of information about each test. To open a new database layout/page with all the curated details for any particular test, click the blue “See” button for that test. The main part of the Test page is a three-tabbed panel. The first tab is open by default and shows data about the laboratory and URLs for the FDA documents. The original EUA Summary and other documents are available for viewing either within the document viewer, e.g. by clicking on “Show EUA Summary”, or by opening a separate browser window outside of the database, e.g. by clicking on the adjacent blue world-wide web buttons.

**Figure 4.**
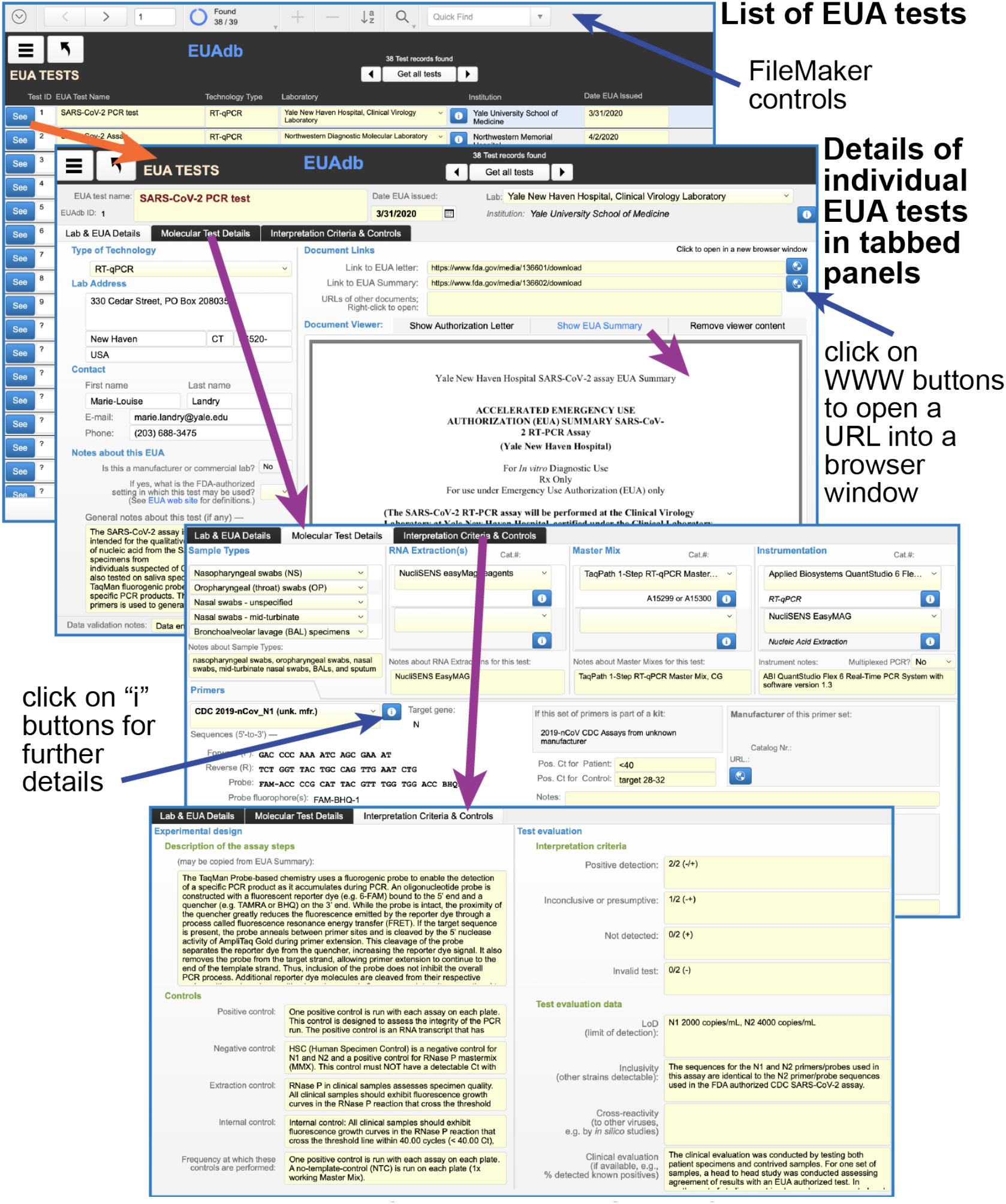
Example showing details for a particular test. See text for explanation.

The second tab (opened by clicking on the “Molecular Test Details” tab title, Fig. 4) shows the sample types, nucleic acid extraction and PCR master mix reagents, instrumentation and primers used for a particular test. These data are shown as rows in “portals” linked to the join tables for those items. Additional information specific to an item may be obtained by clicking on an adjacent “i” button.

The third tab contains information about experimental controls and criteria for interpreting the test data.

Keyword-based queries for tests typically use the list of tests layout to show all the test records in the found set. However, if there is only one such record, the layout automatically switches to the test details layout.

## Findings

Queries of EUAdb reveal several interesting aspects about the diversity of the tests. A plurality—but not a majority— of the tests use the primers originally recommended by the CDC (US Centers for Disease Control and Prevention). The CDC primers target the *N1*, *N2*, and *N3* genes. Other SARS-CoV-2 regions that are targeted include *orf-1ab* and the genes encoding the E, RdRP, and S proteins. Some of the tests only use a single primer pair to detect viral genes, whereas most use at least two and sometimes three. The RT-qPCR result needed to declare a positive result for COVID-19 infection varies. For example, some tests with three primer pairs only require one amplification reaction to declare positivity, while others require at least two out of the three. The controls and sampling methods used also vary widely with the nasopharyngeal swab as the only universal sampling method. Collecting nasopharyngeal samples is invasive and uncomfortable for the patient and might put health care workers at a higher risk of disease transmission^4^. These facts, in addition to recent swab and personal protective equipment shortages, have led to calls for the development of non-invasive, self-sampling methods using saliva^4^. As of this writing, only three tests can utilize saliva specimens (Figure 1), with the Rutgers' test being the only one for which patients can self-collect without health-care worker supervision. Reagents used for RNA extraction and RT-qPCR vary substantially, as does instrumentation (although a plurality of tests use Applied Biosystems QuantStudio for qPCR). As detailed in a separate, co-submitted study from the C. Mason lab (Cornell Univ. Weill Medical School), the LOD (limit of detection) varies by several orders of magnitude across tests. Finally, we found that information provided by the FDA is incomplete with regard to several EUA applications announced prior to March 31, 2020. Of these, six are listed in separate tables on what appears to be a semi-redundant FDA web site^5^. We conclude that at least 12 pre-March 31 EUA applications are not listed on the FDA’s web site; we are contacting the owners of these EUAs to encourage them to provide their EUA submission materials for databasing in EUAdb.

## Conclusion

By providing standardized data and controlled ontology, EUAdb should provide a useful and user-friendly resource for browsing, querying and comparing the different kinds of COVID-19 tests for which the FDA has issued EUAs^2^. The source files can be provided to those who wish to modify the database for their own research purposes.

## Acknowledgements

This work was supported by NIH Fellowship 1F32GM136170 to AW, NSF grant IOS 1656736 to DHAF and an award from the Pershing Square Foundation to JDB. JDB is a founder and director of the following: Neochromosome, Inc., the Center of Excellence for Engineering Biology, and CDI Labs, Inc. and serves on the Scientific Advisory Board of the following: Sangamo, Inc., Modern Meadow, Inc., and Sample6, Inc.

